# Genomic insights into the evolutionary relationships and demographic history of kiwi

**DOI:** 10.1101/2022.03.21.485235

**Authors:** Michael V Westbury, Binia De Cahsan, Lara D Shepherd, Richard N Holdaway, David A Duchene, Eline D Lorenzen

## Abstract

Kiwi are a unique and emblematic group of birds endemic to New Zealand. Deep-time evolutionary relationships among the five extant kiwi species have been difficult to resolve, in part due to the absence of pre-Quaternary fossils to inform speciation events among them. Here, we utilise single representative nuclear genomes of all extant kiwi species (great spotted kiwi, little spotted kiwi, Okarito brown kiwi, North Island brown kiwi, and southern brown kiwi) and investigate their evolutionary histories with phylogenomic, genetic diversity, and deep-time (past million years) demographic analyses. We uncover low levels of gene-tree phylogenetic discordance across the genomes, suggesting clear distinction between species. However, we also find indications of post-divergence gene flow, concordant with recent reports of interspecific hybrids. The four species with available reference assemblies show relatively low levels of genetic diversity, which we suggest reflects a combination of older environmental as well as more recent anthropogenic influence. In addition, we uncover similarities and differences in the demographic histories of the five kiwi species over the past million years, and suggest hypotheses regarding the impact of known past environmental events, such as volcanic eruptions and glacial periods, on the evolutionary history of the group.

## Introduction

New Zealand’s unique and emblematic kiwi, comprising five extant species, represent a highly divergent avian lineage within Palaeognathae. Kiwi are the only members of the Apterygidae family, and are endemic to New Zealand. The species display a number of unusual biological attributes more commonly associated with small mammals. These include a low metabolic rate, lack of colour vision, flightlessness, longevity, and nocturnality [1,2].

The evolutionary origins of kiwi are yet to be fully elucidated. Kiwi are estimated to have diverged from their nearest known relative, the elephant birds (Aepyornithidae), ∼50 million years ago (Ma) [3]. Elephant birds are only known from Madagascar, and it is therefore thought kiwi arrived in New Zealand via flight rather than Gondwanan continental movement and vicariance [4]. Despite such a deep divergence from its closest known relative, the oldest reported kiwi fossil has been dated to at least 19–16 Ma [5]. In contrast, molecular data have suggested the root of living kiwi can be traced back to between ∼15 Ma [6] and ∼5 Ma [7]. Thus, wide uncertainty remains with regard to the lineage leading to extant kiwi, after Apterygidae diverged from Aepyornithidae. Furthermore, deep evolutionary relationships are obscured by the absence of pre-Quaternary fossils to inform speciation events among the extant species.

The five recognized species of kiwi all belong to the genus *Apteryx*. The species are placed into two morphologically and genetically distinct clades [7,8] (Fig. 1). One clade comprises two species, great spotted kiwi (*A. maxima*, formerly *A. haastii* [9]) and little spotted kiwi (*A. owenii*). The other clade includes Okarito brown kiwi (*A. rowi*, also known as rowi), southern brown kiwi (*A. australis*, also known as tokoeka), and North Island brown kiwi (*A. mantelli*). However, despite the deep divergences of these lineages [3,6,7], interspecific hybridization between species from different clades has recently been reported [9].

**Figure 1:**
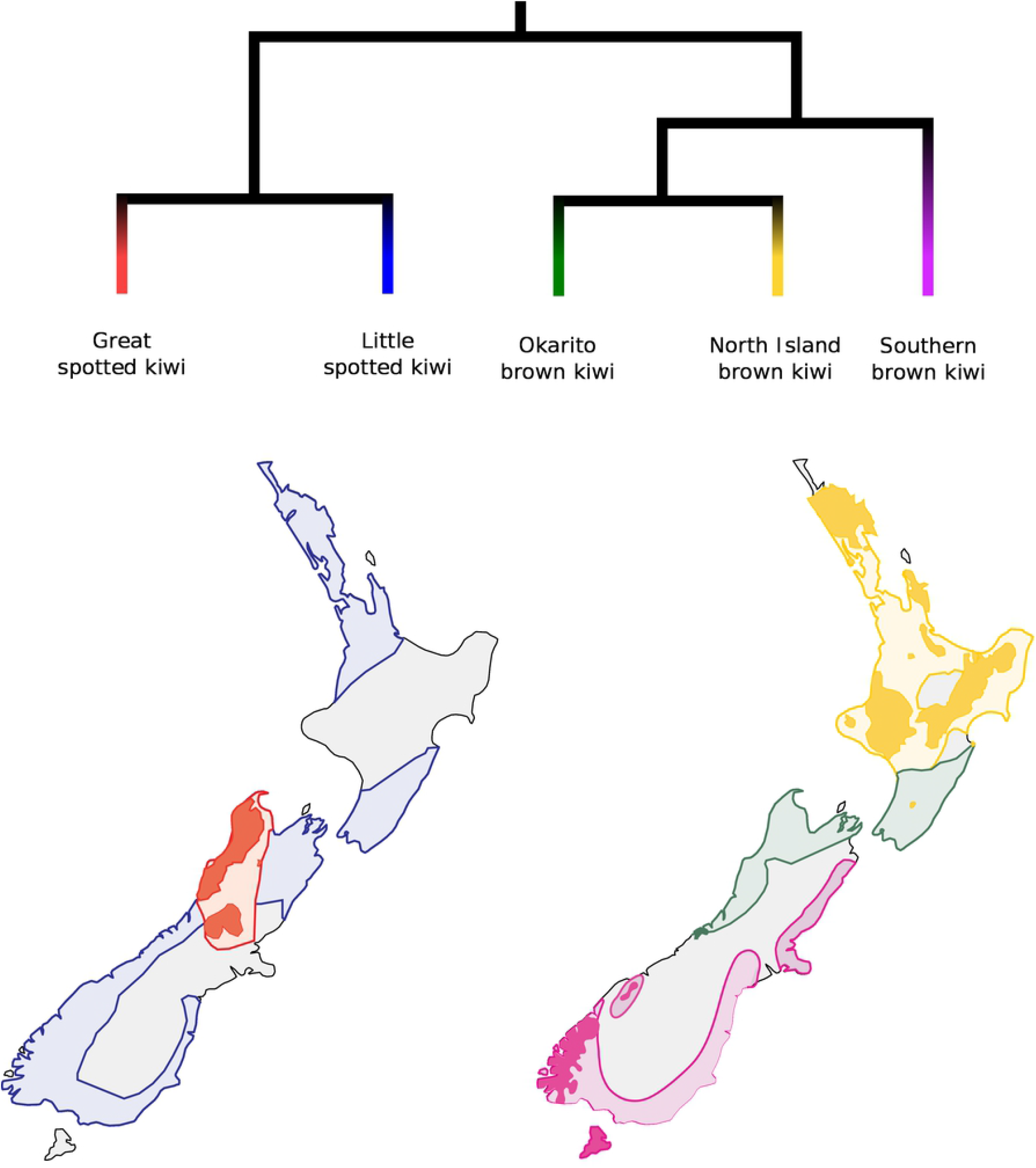
Phylogenetic relationships and distribution ranges of the five extant kiwi species. Distribution ranges are based on [41]. Lightly shaded areas show estimated distributions prior to human arrival and filled in shading shows current distribution. Current distribution for the little spotted kiwi not shown, as it is currently only found on small offshore islands and wildlife sanctuaries. Colours correspond to each respective species.

Prior to human arrival 1,000-800 years ago, kiwi were found from coastal to subalpine habitats across New Zealand, but were most common in lowland rainforest habitats [7]. Partly due to their unique traits, such as flightlessness and longevity, kiwi are highly vulnerable to mammalian predators. Since human colonisation, the distribution range of all five extant kiwi species has been greatly reduced (Fig. 1). The conservation status of four out of the five species is relatively critical (Table 1), and is attributed to population declines following human colonisation [10,11].

**Table 1:** Estimated population sizes and conservation statuses of the five extant kiwi species. DOC - Department of conservation of New Zealand. IUCN - International Union of Conservation of Nature. For southern brown kiwi, DOC conservation statuses vary for different populations, and include nationally critical, nationally vulnerable, and nationally endangered. Population size estimates taken from [41].

| Species | Estimated population size | DOC status | IUCN status |
| --- | --- | --- | --- |
| Great spotted kiwi | 14,000 | Nationally vulnerable | Vulnerable |
| Little spotted kiwi | 1,900 | At risk - recovering | Near threatened |
| Okarito brown kiwi | 600 | Nationally vulnerable | Vulnerable |
| Southern brown kiwi | 24,850 | Population dependent | Vulnerable |
| North Island brown kiwi | 25,100 | At risk -declining | Vulnerable |

Previous studies using short fragments of mitochondrial DNA retrieved from prehistoric and contemporary specimens revealed the genetic consequences of these past range contractions, and suggested significant loss of genetic diversity after the arrival of humans, especially in the Okarito brown, southern brown, and little spotted kiwi [11,12]. In fact, all extant little spotted kiwi are believed to descend from five founding individuals that were placed on the predator-free Kapiti Island in 1912 [13].

The use of genome-wide data can greatly enhance our existing understanding of kiwi relationships, as they allow analyses of species deep-time evolutionary histories. *De novo* nuclear genome assemblies are available from four of the five kiwi species (great spotted kiwi, little spotted kiwi, Okarito brown kiwi, North Island brown kiwi) [2,14], and resequencing data are available from the southern brown kiwi [15]. Here, we utilise single genome representatives of each species to better understand the interspecific evolutionary relationships among kiwi species, and to elucidate how past environmental changes (up to 1 Ma) may have impacted their demographic histories.

## Materials and Methods

### Data

We downloaded raw Illumina sequencing reads and genome assemblies from each of

the four available kiwi species from Genbank: great spotted kiwi, little spotted kiwi, Okarito brown kiwi, and North Island brown kiwi. We downloaded the raw reads for a southern brown kiwi from the individual with the highest number of available reads. To act as an outgroup for the phylogenomic analyses, we downloaded raw sequencing reads and the genome assembly of emu (*Dromaius novaehollandiae*). For estimates of comparative genetic diversity, we downloaded raw sequencing reads and the genome assemblies of three additional Paleognath species: ostrich (*Struthio camelus*), southern cassowary (*Casuarius casuarius*), and greater rhea (*Rhea americana*). Accession code details for all species can be found in supplementary table S1.

### Data processing

We trimmed adapter sequences and removed reads shorter than 30 bp from the downloaded raw reads using skewer [16], mapped the trimmed reads to the specified reference genome assembly using BWA v0.7.15 [17] and the mem algorithm. We parsed the output and removed duplicates with SAMtools v1.6 [18]. Furthermore, as some downstream analyses required the removal of sex chromosomes, we independently determined which scaffolds were most likely autosomal in origin for each reference genome assembly used in the current study. We found putative sex chromosome scaffolds by aligning each reference genome assembly to the chicken (*Gallus gallus*) Z and W chromosomes (Genbank accessions: Z - CM000122.5, W - CM000121.5). Alignments were performed using satsuma synteny [19] and default parameters.

### Phylogenomics

We performed multiple phylogenomic analyses using different mapping reference genome assemblies, to assess the robustness of our results to reference choice. We independently mapped the raw reads of all five kiwi species to two reference genome assemblies: emu and great spotted kiwi. We also mapped the emu raw reads to the emu reference genome assembly. From the resultant mapped files, we built majority rules consensus sequences (-doFasta 2) in ANGSD v0.921 with the following parameters; - mininddepth 5 -minmapq 30 -minq 30 -uniqueonly 1, and only included autosomal scaffolds >100kb in length (-rf). We extracted 20 kb windows with a 1 Mb slide from each consensus sequence using bedtools v2.26.0 [20]. We performed maximum likelihood phylogenetic inference for each window in IQ-TREE2 [21], using the best GTR+R+F model according to the Bayesian information criterion [22]. Using the estimated gene trees, we inferred the species tree of kiwi under each reference genome assembly, using the summary multispecies coalescent as implemented in ASTRAL v4.10 [23]. The species tree inferred for each of the two datasets was used as the focal tree to calculate gene- and site-concordance factors for each branch in IQ-TREE2 [24].

### Quantifying introgression via branch lengths

To investigate whether phylogenetic discordance among all possible kiwi triplets [[A,B],C] can be explained by incomplete lineage sorting (ILS) alone, or by a combination of ILS and gene flow, we implemented Quantifying Introgression via Branch Lengths (QuIBL) [25] on the dataset obtained when mapping the kiwi and emu raw read data to the emu reference genome. We used the kiwi and emu mapped to the emu dataset due to the requirement of an outgroup. Furthermore, since this analysis relies on the relative ages of nodes, we only used loci with nearly constant evolutionary rates among lineages. Specifically, we rooted each gene tree with the emu and excluded those loci with a coefficient of variation in root-to-tip length >0.01. We ran QuIBL specifying the emu as the overall outgroup (totaloutgroup), to test either ILS or ILS with gene flow (numdistributions 2), the number of total EM steps as 50 (numsteps), and a likelihood threshold of 0.01. We determined the significance of gene flow by comparing values of BIC1 (ILS alone) and BIC2 (ILS and gene flow). If the difference between BIC1 and BIC2 was greater than 10, we assumed incongruent topologies arose due to both ILS and gene flow. With a difference of less than 10, we assumed ILS alone.

### End of lineage sorting/gene flow

To estimate the point in time when the genomes of the five kiwi species had fully coalesced, putatively indicating an end of lineage sorting and/or gene flow, we used the F1 hybrid Pairwise Sequentially Markovian Coalescent model, hPSMC [26]. Previous studies have shown that the results of hPSMC are not significantly influenced by mapping reference [27,28], and we therefore performed each analysis once, with the raw reads from the five kiwi species mapped to the great spotted kiwi reference genome assembly. We constructed haploid consensus sequences for each of the kiwi individuals using the same consensus sequences as the phylogenomic analysis. We merged the resultant haploid consensus sequences pairwise into a pseudo diploid sequence using a python script available as part of the hPSMC toolsuite. The resultant pseudodiploid sequences were run through a Pairwise Sequentially Markovian Coalescent model (PSMC) [29].

To calibrate the PSMC plots, we calculated an *Apteryx* average mutation rate. We did this by first calculating the average pairwise distance for each species pair (Supplementary table S2), and dividing that by 2 * the previously published divergence times [3]. We calculated the pairwise distances in ANGSD v0.921 [30] using a consensus base call approach, with all species mapped to the great spotted kiwi reference genome assembly, and applying the same filters as for the phylogenomic analyses with the additions of only including sites found in all individuals (-minInd 5), and print a distance matrix (-makematrix 1). This resulted in a mutation rate of 8.0×10^−10^ per year or 2.0×10^−8^ per generation, assuming a generation time of 25 years [7].

From the PSMC output, we manually estimated the pre-divergence Ne of each pseudodiploid genome by outputting the text file (-R) using the plot script from the PSMC toolsuite. Using the pre-divergence Ne estimated from this output, we ran simulations to infer the intervals during which the pseudodiploid genomes coalesce between each species pair using ms [31]. Simulation commands in ms were automatically produced with the hPSMC_quantify_split_time.py python script from the hPSMC toolsuite, while specifying the pre-divergence Ne and the time windows we wanted to simulate, and the remaining parameters as default. The time intervals and pre-divergence Ne for each species pair can be found in supplementary table S3. We plotted the results and found the simulations with an exponential increase in Ne to be closest to the empirical data, between 1.5-fold and 10-fold the value of the pre-divergence Ne. The divergence times from these simulations were taken as the time interval during which lineage sorting and/or gene flow stopped. We considered the portion between 1.5-fold and 10-fold the value of the pre-divergence Ne, as suggested in previous work [26]. This was done to capture the portion of the hPSMC plot most influenced by the divergence event. The lower bound is set to control for pre-divergence increases in population size, and the upper bound is to avoid exploring parameter space in which little information is present.

### Autosome-wide heterozygosity and inbreeding estimates

We calculated autosome-wide levels of heterozygosity and runs of homozygosity (ROH) for each species using the software ROHan [32]. The raw reads of the four species for which conspecific reference genome assemblies were available, were independently mapped to each said assembly. Raw reads of the southern brown kiwi, which does not have an available conspecific assembly, were mapped to the North Island brown kiwi assembly. We ran ROHan three times independently, using different window sizes (500 kb, 1 Mb, 2 Mb), while keeping the other parameters as default. The default parameters specify a window as being a ROH if it has an average heterozygosity of less than 1×10^−5^. We calculated autosome-wide heterozygosity for the non-kiwi palaeognath species (emu, ostrich, southern cassowary, greater rhea) in the same way, but using only a 1 Mb window size.

### Demographic reconstruction

We ran demographic analysis on diploid consensus genomes from each kiwi species, each mapped to their conspecific reference genome assembly and the southern brown kiwi mapped to the North Island brown kiwi, using PSMC [29]. We called diploid genome sequences using SAMtools and BCFtools v1.6 [33], specifying a minimum quality score of 20 and minimum coverage of 10. However, we also created a diploid consensus genome with Okarito brown kiwi mapped to the North Island brown kiwi, as it has previously been shown that mapping to the Okarito brown kiwi reference genome assembly may be problematic for PSMC analysis [34]. As Prasad et al. did not assess the reliability of the little spotted kiwi as mapping reference, we further assessed the reliability of the PSMC results of this species by mapping to the conspecific reference genome assembly, as well as to the great spotted kiwi and North Island brown kiwi reference genome assemblies. Before running PSMC, we removed scaffolds found to align to sex chromosomes in the previous step, and removed scaffolds shorter than 100 kb. We ran PSMC specifying standard atomic intervals (4+25*2+4+6), and performed 100 bootstrap replicates to investigate support for the resultant demography. We plotted the output using a mutation rate of 2.0×10^−8^ per generation, assuming a generation time of 25 years as detailed above for the hPSMC analysis.

## Results

### Interspecific evolutionary relationships

Regardless of the genome assembly selected as mapping reference, the vast majority of windows in our phylogenomic analysis support the grouping of little spotted kiwi and great spotted kiwi as sister species, and the grouping of Okarito brown kiwi, North Island brown kiwi, and southern brown kiwi in a separate clade, with the southern brown kiwi being the earliest to diverge within the brown kiwi clade (Fig 1, Supplementary tables S4). However, within the brown kiwi, alternative gene tree topologies occurred at relatively high frequency (up to 11%). Site concordance factors also found high levels of discordance within this clade (up to 26%). Further examination using QuIBL to assess if alternative topologies may have arisen due to ILS alone, or to both ILS and gene flow, indicated the few topological discordances most likely represent ILS rather than gene flow (Supplementary table S5).

To further investigate signs of post-divergence gene flow between kiwi lineages, we ran the F1 hPSMC described above. We find that, despite the deep divergences of kiwi lineages, their genomes coalesce relatively recently. Between the two major clades, we found that lineage sorting and/or gene flow ceased 2.6-1.8 Ma. This pattern was the same, regardless of which individuals were used in the pairwise comparison (Supplementary fig. S1). Within clades, we find lineage sorting was complete and/or gene flow ceased between great spotted kiwi and little spotted kiwi 700-400 thousand years ago (kya) (Supplementary fig. S2), and similarly between North Island brown kiwi and Okarito brown kiwi 800-500 kya (Supplementary fig. S3). Lineage sorting was complete and/or gene flow ceased 1.2 Ma-800 kya between southern brown kiwi and both the North Island brown kiwi and the Okarito brown kiwi (Supplementary fig. S4).

### Autosome-wide heterozygosity and inbreeding estimates

We find the highest levels of autosome-wide heterozygosity in North Island brown kiwi, followed by great spotted kiwi, southern brown kiwi, Okarito brown kiwi, and little spotted kiwi (Fig 2A). To further contextualise these estimates, we calculated the autosome-wide heterozygosity estimates of four other palaeognaths (emu, ostrich, southern cassowary, greater rhea). Our analysis revealed that kiwi species have relatively low levels of heterozygosity compared to the other palaeognaths, except for the southern cassowary, whose heterozygosity only exceeded that of little spotted kiwi.

**Figure 2:**
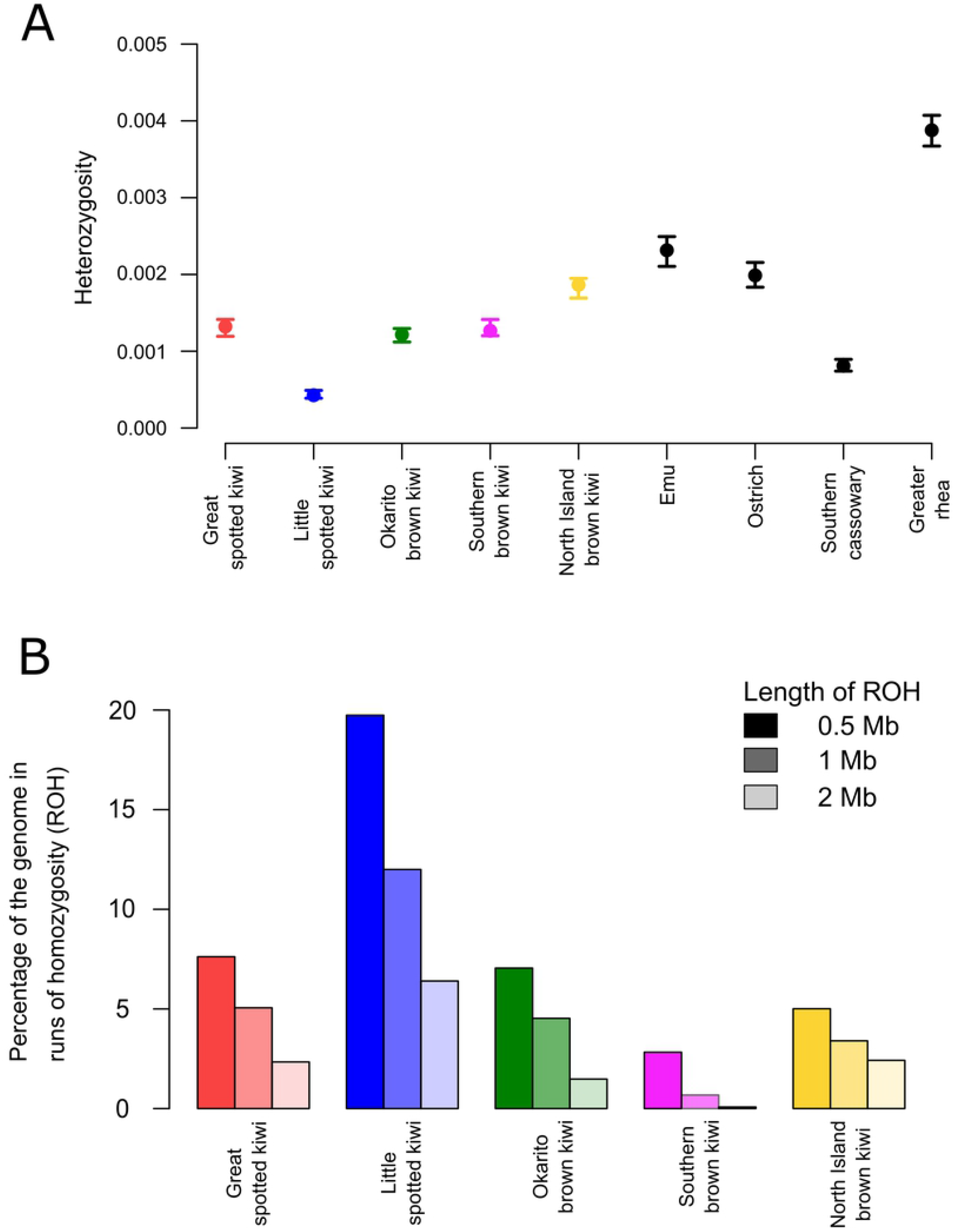
Genetic diversity and inbreeding estimates. (A) Autosome-wide heterozygosity calculated using ROHan for the four kiwi species with conspecific reference genomes, and four other palaeognath species. Error bars show maximum and minimum values. (B) Autosomal runs of homozygosity calculated using various window sizes for the four kiwi species with conspecific reference genomes. Differential shading shows the window size used.

We further investigated whether recent inbreeding may have caused the low levels of autosome-wide heterozygosity using runs of homozygosity (ROH) analyses. As expected, the number of ROH increased when specifying increasingly smaller window sizes as a ROH (Fig 2B). Moreover, the general trend reflected what would be expected based on autosome-wide heterozygosity levels. That is, the species with the lowest overall heterozygosity (little spotted kiwi) also had the most ROH, and the species with the highest overall heterozygosity (North Island brown kiwi) had the least ROH. We found the lowest levels of ROH in the southern brown kiwi (Fig 2B).

This pattern was not as obvious in the other palaeognath species. The individual with the highest levels of diversity (rhea) also showed considerable levels of ROH (1 Mb window size; 2.2% of the genome in ROH). However, the individual with the lowest diversity (southern cassowary), also had the highest levels of ROH (1 Mb window size; 8.42% of the genome in ROH). The remaining two species had low levels of ROH: emu - 0.50% and ostrich - 0.64%.

### Intraspecific demographic histories

Our assessment of how mapping reference influences PSMC results of little spotted kiwi showed similar results to those reported previously for Okarito brown kiwi [34]; we find relatively consistent results when mapping the little spotted kiwi raw reads to both North Island brown kiwi and to great spotted kiwi, but much reduced effective population size (Ne) values when mapping to little spotted kiwi (Supplementary fig. S5). Therefore, we based our inferences on the results retrieved when mapping to its closest relative (great spotted kiwi).

We find Ne in North Island brown kiwi was relatively high and stable over the past 1 Ma, with several fluctuations within the past 100-50 kya (Fig. 3). However, the bootstrap support values during the latter time period show considerable uncertainty, suggesting little confidence in these fluctuations. The southern brown kiwi also exhibited relatively stable Ne values until ∼200 kya where there was a continual decrease in Ne until present. Ne in Okarito brown kiwi decreased 1-0.5 Ma, followed by a plateau and slight increase, then a more rapid decline in the past ∼100 kya. The two remaining species, little spotted kiwi and great spotted kiwi, exhibit similar demographic trends. Both species show a continuous gradual decline in Ne over the past 1 Ma.

**Figure 3:**
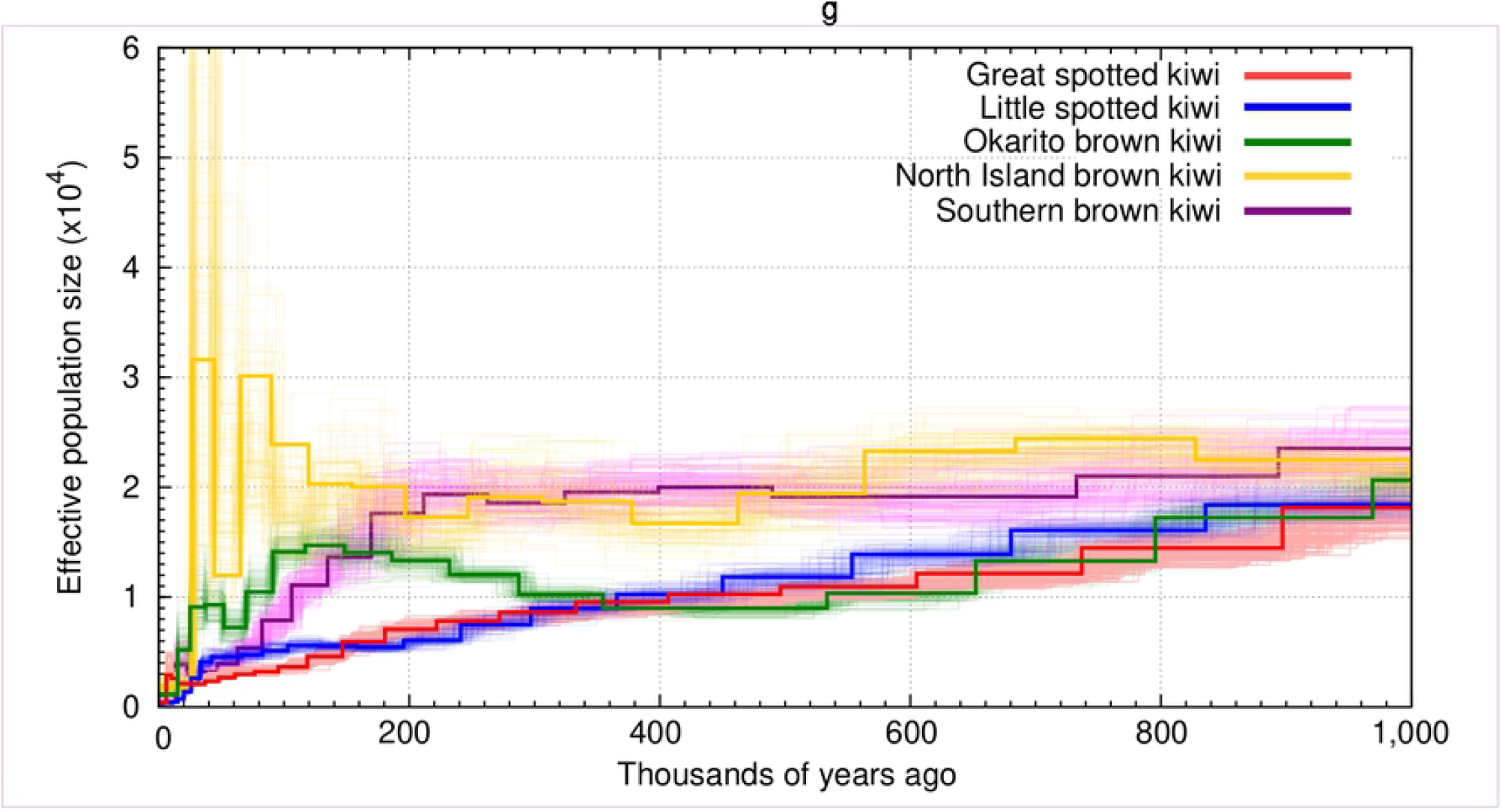
Effective population sizes over the past one million years of the five extant kiwi species calculated using PSMC. Faded lines show the 100 bootstrap replicates produced for each species.

## Discussion

We recovered phylogenetic relationships among the five kiwi species consistent with previous research using mtDNA sequences and nuclear data [3,7,15,35](Fig. 1).We found these groupings to be highly conserved across windows; the few phylogenetic discordances observed across the genomes most likely arose due to ILS, as opposed to gene flow (Supplementary table S5). However, despite the clear differentiation between species when considering gene (window) tree concordance factors, our hPSMC results support continual gene flow between all species long after they initially diverged (Figs S1-3). This is more in line with the high levels of discordance at the site level (Supplementary table S6), especially within the Okarito, southern, North Island brown kiwi clade. This scenario is more congruent with recent reports of hybridisation between kiwi sister species, and between kiwi species from each of the two distinct clades [9]. Although all species now exhibit relatively restricted ranges (Fig. 1), fossil evidence indicates broader ranges, at least throughout the Holocene [7]

The presence of interspecific gene flow between kiwi species may explain the large discrepancies between molecular estimates of their divergence times. Estimates of the stem divergence of the five extant species have differed markedly. These include, but are not limited to, ∼14.5 Ma [6], ∼13.4 Ma [36], ∼12.31 Ma [3], ∼11.3 Ma [35], ∼8.5 Ma [4,37], and ∼5.9 Ma [7]. Even when considering the most recent split (∼5.9 Ma), our hPSMC results suggest the genomes completely coalesced between 2.6 Ma and 1.8 Ma, which may be interpreted as the continuation of lineage sorting for a long time post divergence, or continued gene flow.

Interspecific gene flow is well documented in avian species [38]. However, this is generally correlated with their high dispersal rates through flight. Although kiwi are known to have evolved flightlessness independently from other palaeognaths after their arrival to New Zealand [3,6], a lack of fossils makes it difficult to reliably determine when flightlessness evolved. Nevertheless, based on a recent fossil find, kiwi are thought to be flightless from at least the Middle Pleistocene (∼1 Ma) [39]. A lack of migration through flight makes the ability for continued hybridisation more profound, and can only be fully addressed with genomic data from more individuals sampled from across their respective ranges.

Our finding of very low diversity in little spotted kiwi is congruent with previous studies based on microsatellite data [40] and population-level nuclear genomes [15]. Our finding of low diversity in this species — at least as represented by our sampled individual — may reflect the high levels of inbreeding we uncovered, putatively due to the recent history of this species. In 1912, five little spotted kiwi were moved to an offshore, predator-free island (Kapiti Island), and their descendants have been subsequently translocated to other offshore islands and predator-free wildlife sanctuaries. Therefore, the entire population of the species, estimated at approximately 2,000 individuals [41], is descended from at most five founders; a recent microsatellite study indicated perhaps only three founder individuals [12]. Such a low number of founders would undoubtedly lead to high levels of inbreeding. However, this is in contrast to a lack of signs of inbreeding suggested by a recent population genomic study [15]; we suggest this may be a byproduct of the computational approach employed in the study, rather than a biological signal. Mapping to a phylogenetically distant reference has been shown to essentially remove signs of inbreeding [34]. Bemmels et al (2021) did not map the little spotted kiwi to a conspecific reference genome, and therefore the lack of inbreeding may be the result of mapping biases.

Okarito brown kiwi experienced a bottleneck to ∼150 birds in the 1990s [42]. However, we find similar diversity and inbreeding levels to great spotted kiwi (Fig 2). Although the range of great spotted kiwi is thought to have become highly fragmented and reduced by 30% since human arrival in New Zealand [43], an estimated 14,000 great spotted kiwi remain [41]. Based on 11 microsatellite loci, it was found that great spotted kiwi had high levels of genetic diversity compared with other kiwi species, and showed no evidence of a recent bottleneck (within the last 100 generations) [44]. Therefore, if genetic diversity alone predicts population size, great spotted kiwi should display higher levels of diversity than Okarito brown kiwi. However, as our Okarito brown kiwi individual was sampled in 1993 (and may have hatched as early as the 1940s), our findings likely do not reflect the genetic impact of the recent bottleneck in the 1990s. This suggests that the Okarito brown kiwi population at Okarito, from which this sample arose, likely had higher levels of diversity pre-bottleneck, and exemplifies the problems when analysing genomes from individuals sampled prior to bottleneck events. Finally, our finding of North Island brown kiwi having the highest levels of diversity is consistent with higher census population size (25,000) than the other species [41].

Although recent events can be used to explain low levels of diversity in kiwi species, the longevity and generation times (longevity up to 45 years and 25 year generation time [45]) of kiwi means the genomic diversity of modern individuals may not accurately reflect recent demographic events (order of 100s of years). By investigating changes in Ne over evolutionary time scales, we observe all species have reduced in Ne within the last 40-20 kya, and in some cases much longer time periods, suggesting low genetic diversity is not only a recent anthropogenically-driven phenomenon. However, the ROH we uncovered in all species with a conspecific reference genome (great spotted kiwi, little spotted kiwi, Okarito brown kiwi, North Island brown kiwi) (Fig 2) suggests recent bottlenecks and inbreeding may have further exacerbated already low levels of diversity in kiwi.

Owing to a lack of fossil evidence, it is difficult to know how kiwi populations responded to past climatic and environmental events. However, genomic data allows for the exploration of changes in population size over evolutionary timescales, and to suggest hypotheses based on correlations with known environmental events.

We observe relatively stable Ne in the North Island brown kiwi over the past one million years. During this period, the central North Island experienced several major rhyolitic eruptions, as evidenced by tephras in ocean cores to the east and northeast of the North Island [46]. The ash from these eruptions rarely reached the northern peninsula of the North Island, providing a putative refuge for this population. Therefore, the northern distribution of North Island brown kiwi (Fig. 1) may have facilitated this long-term stability in population size, by providing a relatively constant environment. Similarly, we also observe a stable Ne in the southern brown kiwi 1 Ma - 100 kya. The highly southern range of this species may also have allowed it to persist at relatively stable population sizes, despite eruptions.

In contrast, both great and little spotted kiwi show a general trend of declining Ne throughout the past one million years. As the genetic data of both taxa are from the South Island, these results may only represent changes to South Island populations, and therefore climatic and other events affecting only the South Island may have driven the declines. Both volcanic fallout and repeated glaciations – or repeated *de*glaciations – may have had long-term negative effects on the populations of both kiwi taxa. However, little spotted kiwi ranged on both the North and South Islands pre-human colonisation (Fig 1), so the inclusion of genomic information from extirpated North Island populations may provide more insights into whether these declines were species-wide or local occurrences. There are few museum specimens of little spotted kiwi from the North Island, with only two live birds collected, both in the 19th century [8]. Despite the rarity of museum specimens, Holocene bones of little spotted kiwi may also provide a means to study species-wide demographic patterns, although DNA preservation from some sites, such as dunes, may be poor.

The Ne of Okarito brown kiwi also shows a decreasing trend, suggestive of similar mechanisms to those in the great and little spotted kiwi. However, the trajectory deviates with a plateau in Ne, before rapidly decreasing ∼100 kya. Associated with this decline is the end of the most recent interglacial (prior to the Holocene), Isotope Stage 5e (130 - 116 kya [47]). A change in Ne at this time could be caused by the restriction of population movement or a reduction in population size, both of which could have been a consequence of the environmental instability of this period. However, Okarito brown kiwi has experienced many glacial and interglacial periods before this event, and therefore it is difficult to determine coincidence or causation.

All five kiwi species display a rapid and steep decline in Ne coinciding with the most recent global volcanic super eruption (Fig. 3). The Oruanui eruption of Taupo volcano 25.6 kya deposited 1000 km^3^ of tephra across New Zealand, from Auckland in the North Island to the Waitaki River in the South Island [48,49]. Such a catastrophic event is expected to have destroyed most of the New Zealand biota between 38° and 45° South. If the decline in Ne in association with this major eruption was exhibited by just one taxon, then little case could be made for cause and effect. However, the analogous temporal pattern of a population decline across all five species lend strong support for a significant environmental driver.

Although it is difficult to precisely pinpoint the driver of past population fluctuations, we form a number of hypotheses mostly based on correlations with volcanic activity. To date, few studies have used whole-genome data to investigate the longer-term demographic histories of New Zealand avian species, and those that have, have focussed on glaciation as a major influence [50–52]. As more genomes from a wider range of New Zealand species are made available, it will be possible to test whether volcanic eruptions may have also been a previously overlooked major evolutionary force shaping the biota of New Zealand.

## Conclusions

By analysing publicly available genomic data, we uncover genomic indications of post-divergence gene flow among all kiwi species, a factor that may have led to difficulties in accurately identifying species divergence times in previous studies. Moreover, we gained novel insights into the demographic history of kiwi species over the past one million years, the timing of which is associated with various environmental factors. There has been limited research on the impact of volcanic events on New Zealand avifauna; our study provides evidence that these environmental disruptions may have played a bigger role than previously thought, opening new avenues of future research.

## Acknowledgements

This work was supported by the Independent Research Fund Denmark | Natural Sciences, Forskningsprojekt 1, grant no. 8021-00218B, and the Villum Fonden Young Investigator Programme, grant no. 13151, to EDL. We would like to acknowledge the Te Parawhau Trust and Waikaremoana iwi, Te Runanga o Ngai Tahu, Te Ātiawa Manawhenua Ki Te Tau Ihu Trust, who provided guidance to the authors who generated the kiwi genomic data, on which our study is based. The authors declare no conflicts of interest.

## Author contributions

Conceptualization, MVW; Formal analysis, MVW, BDC, DAD; Writing – Original Draft, MVW; Writing – Review & Editing, MVW, LDS, RNH, EDL; Writing – Final revisions, all authors; Funding Acquisition, EDL; Supervision, EDL, MVW.

## Data availability

All raw data used in this manuscript were previously published and the accession codes are listed in supplementary table S1.

